# Motor plans under uncertainty reflect a trade-off between maximizing reward and success

**DOI:** 10.1101/2021.11.21.469448

**Authors:** Aaron L. Wong, Audrey L. Green, Mitchell W. Isaacs

## Abstract

When faced with multiple potential movement options, individuals either reach directly to one of the options, or initiate a reach intermediate between the options. It remains unclear why people generate these two types of behaviors. Using the go-before-you-know task (commonly used to study behavior under choice uncertainty), we examined two key questions. First, do these two types of responses reflect distinct movement strategies, or are they simply examples of a more general response to choice uncertainty? If the former, the relative desirability (i.e., weighing the likelihood of successfully hitting the target versus the attainable reward) of the two target options might be computed differently for direct versus intermediate reaches. We showed that indeed, when exogenous reward and success likelihood (i.e., endogenous reward) differ between the two options, direct reaches were more strongly biased by likelihood whereas intermediate movements were more strongly biased by reward. Second, what drives individual differences in how people respond under uncertainty? We found that risk/reward-seeking individuals generated a larger proportion of intermediate reaches and were more sensitive to trial-to-trial changes in reward, suggesting these movements reflect a strategy to maximize reward. In contrast, risk-adverse individuals tended to generate more direct reaches in an attempt to maximize success. Together, these findings suggest that when faced with choice uncertainty, individuals adopt movement strategies consistent with their risk/reward-seeking tendency, preferentially biasing behavior toward exogenous rewards or endogenous success and consequently modulating the relative desirability of the available options.

## INTRODUCTION

When faced with selecting between multiple potential options, we plan our response by carefully weighing all available options in an attempt to maximize our performance. When individuals are required to begin their movement before the correct option can be identified, individuals often respond in one of two distinct ways. First, individuals may commit to one of the options before movement onset, and consequently initiate a movement toward that option (i.e., a direct response). In contrast, individuals may generate a movement that lies partway between the available options (i.e., an intermediate response). One popular hypothesis explaining why these intermediate movements are generated proposes that individuals prepare a motor plan to each of the potential options, and that averaging between these plans (unintentional or otherwise) gives rise to intermediate movements (Chapman et al., 2010; Cisek, 2007; Enachescu et al., 2021; Gallivan et al., 2018, 2016). That is, under this averaging hypothesis there is no distinct motor plan to produce an intermediate response; instead, the intermediate “plan” is simply comprised of a weighted average of direct-response plans. In contrast, increasing evidence favors an alternative hypothesis that intermediate responses reflect a strategy to improve performance; such a strategy delays commitment to one option until more information can be attained and minimizes the cost of generating motor corrections (Alhussein and Smith, 2021; Dekleva et al., 2018; Hudson et al., 2007; Nashed et al., 2017; Onagawa and Kudo, 2021; Wong and Haith, 2017). Under this alternative hypothesis, an intermediate movement is therefore deliberately planned as a strategic response to uncertainty.

Aside from deciding whether to generate a direct or an intermediate response, another important decision governing how people respond in the face of uncertain options is to weigh the relative desirability of the two options. In particular, prior research has demonstrated that individuals consider both the potential rewards available in the environment as well as the likelihood with which they will be successful at attaining those rewards (for reviews, see Shadmehr et al., 2019; Wolpert and Landy, 2012). Both when choices are discrete (e.g., direct reaches or saccades) as well as when intermediate movements are generated, movements tend to be biased toward the more rewarding (Chapman et al., 2015; Liston and Stone, 2008; Sugrue et al., 2004) or likely (Chapman et al., 2010; Liston and Stone, 2008) option, reflecting an effort to maximize reward or task success respectively. Note, however, that it is possible to independently manipulate exogenous rewards and the likelihood of success (i.e., endogenous rewards), even in some cases setting these factors in opposition. When both reward and likelihood are unequal between the two options, behavior is ideally determined according to the expected value (i.e., reward times likelihood) of each available option (Johnson and Ratcliff, 2014; Teichert and Ferrera, 2010).

However, neuroeconomics reveals that individuals do not evaluate options according to their objective expected value, but according to their subjective value (utility) according to the degree to which an individual is risk/reward-seeking (Glimcher and Rustichini, 2004; Kahneman and Tversky, 1979; Mongin, 1998). Thus when faced with uncertainty about which of multiple options to choose, individuals may weigh likelihood or reward more heavily in their decisions, leading to preference biases even in cases where the expected values of the options are equal. Specifically, individuals may be biased to reach toward one target more often than the other when making direct reaches, and their initial reach direction may be biased closer to one target than the other for intermediate reaches. Our first goal was therefore to examine whether reward and likelihood contributed equally when determining the relative subjective values of the two reach options – as determined by a bias in reach direction when the objective value of the two targets was matched – and whether this bias was analogous for both direct and intermediate reaches.

Prior studies examining intermediate reaches have all focused their analyses at the group level. What is frequently not discussed is the fact that individuals often differ greatly in how they respond in these tasks, even though every participant is presented with the exact same set of options. While some individuals preferentially respond by generating intermediate responses, others produce only direct responses, and still others produce a mixture of both behaviors. No clear rationale has been proposed as to why, given the exact same set of options on every trial, a mixture of direct and intermediate responses are produced across individuals, let alone why different individuals will respond with different proportions of these two types of behaviors. Our second goal, therefore, was to examine whether differences in the degree to which individuals preferentially generate direct or intermediate movements might be driven by differing degrees of risk/reward sensitivity across individuals.

To answer these questions, we asked participants to perform a go-before-you-know paradigm in which individuals were shown two targets – only one of which was the “correct” target that would yield a reward when hit – and were required to initiate their reach before the correct target was revealed. We varied the reward and likelihood associated with each target such that one target was more rewarding while the other was more likely to be correct, and evaluated the degree to which reward and likelihood bias behavior for both direct and intermediate reaches. We also estimated individual risk/reward sensitivity and examined how that modulated behavior at the individual-subjects level. Together, these findings shed light on how individuals decide to move under uncertainty.

## RESULTS

### Frequency and reward bias behavior to different extents

In experiment 1, participants completed a go-before-you-know task in which participants were presented with two potential targets and were required to initiate a reach before the correct target was revealed (Chapman et al., 2010; Wong and Haith, 2017). We modified this task by assigning each target a particular frequency (likelihood) and reward value (Fig. 1A), with the exact frequency or reward varying across blocks. Following completion of this task, reaches were sorted according to the frequency and reward ratios presented on that trial, and each reach was classified as heading directly to a target or intermediate between the targets (Fig. 1C; see Methods). In general, participants as a group generated a mixture of direct and intermediate reaches in all conditions. Gaussian distributions (bimodal for direct reaches and unimodal for intermediate reaches) were fit to the data to estimate reaching biases exhibited in each condition (Fig. 2). Because not every participant generated both direct and intermediate reaches in every condition, we used bootstrapping to estimate the relationship between the biases induced by varying reward and frequency for direct versus intermediate reaches (see Methods). In general, participants changed their reach biases as the relative frequencies and rewards of the two targets were modulated. For direct reaches, if the frequency of the two targets was equal (Fig. 3A, top panel), participants tended to make proportionally more reaches toward the target with the larger reward. This effect increased as the reward ratio of the two targets increased (slope = 0.01 ± 0.0038 SD). Likewise, if the two targets were rewarded equally (Fig. 3A, left panel), participants tended to make proportionally fewer reaches toward the target that was less frequently revealed to be the correct target (i.e., toward the target that would have been more rewarded if the reward ratio was not 1:1), and this effect increased with the frequency ratio (slope = −0.04 ± 0.0041 SD). For intermediate reaches (Fig. 3B), a similar trend emerged although it was more difficult to detect by eye. Under conditions when the frequency of the two targets was equal (Fig. 3B, top panel), participants tended to exhibit initial reach directions that became increasingly biased toward the more rewarding target as the reward ratio increased (slope = 0.12 ± 0.10). When the rewards assigned to the two targets were equal (Fig. 3B, left panel), participants tended to reach in a direction that was biased toward the more frequent target particularly as the frequency ratio increased (slope = −0.37 ± 0.082).

**Figure 1.**
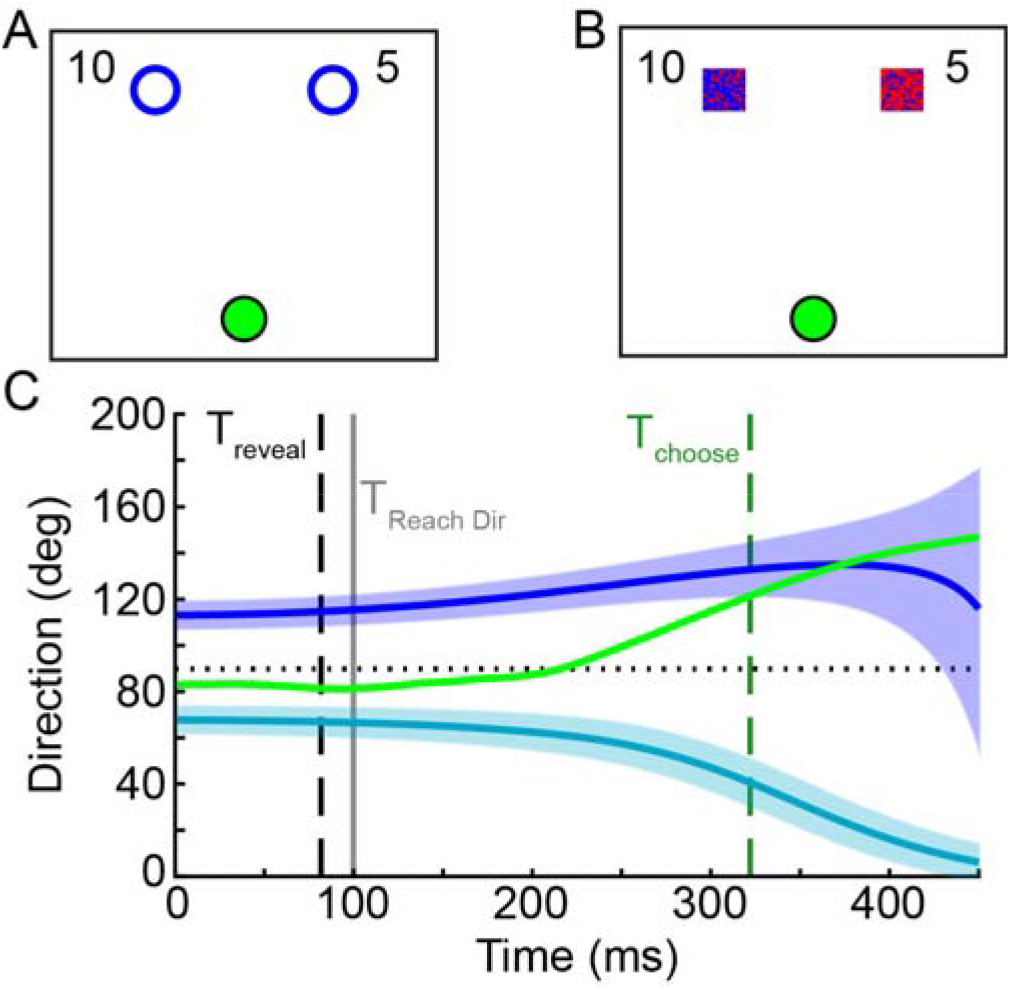
Methods for Experiments 1 and 2. Participants completed a go-before-you-know task in which they observed 2 targets and were required to begin moving; only after they began their movement would the true target would be revealed. (A) In Experiment 1, one of the two target options could occur more frequently, and the other target could result in more reward. Reward offers appeared explicitly on screen (e.g., 10 cents versus 5 cents, or a reward ratio of 2:1); the two targets were otherwise identical and participants had to learn the frequency manipulation through experience. (B) In Experiment 2, the target probability was modulated on each trial instead of as a blockwise frequency; target probability on the current trial was indicated by the color of the target (the degree of redness reflects the probability of being correct, e.g., 80% versus 20%). (C) At each timepoint, the direction of the hand (green) was computed relative to the direction required to be moving toward one of the two targets (blue, teal). The time at which the hand was heading directly toward one of the two target options (T_choose_, dashed green) was compared to the time when the correct target was revealed (T_reveal_, dashed black). If T_choose_ occurred prior to T_reveal_ the movement was classified as direct, otherwise it was classified as intermediate. This was estimated independently of when the initial reach direction was measured, which was always 100 ms into the reach (TReachDir, gray).

**Figure 2.**
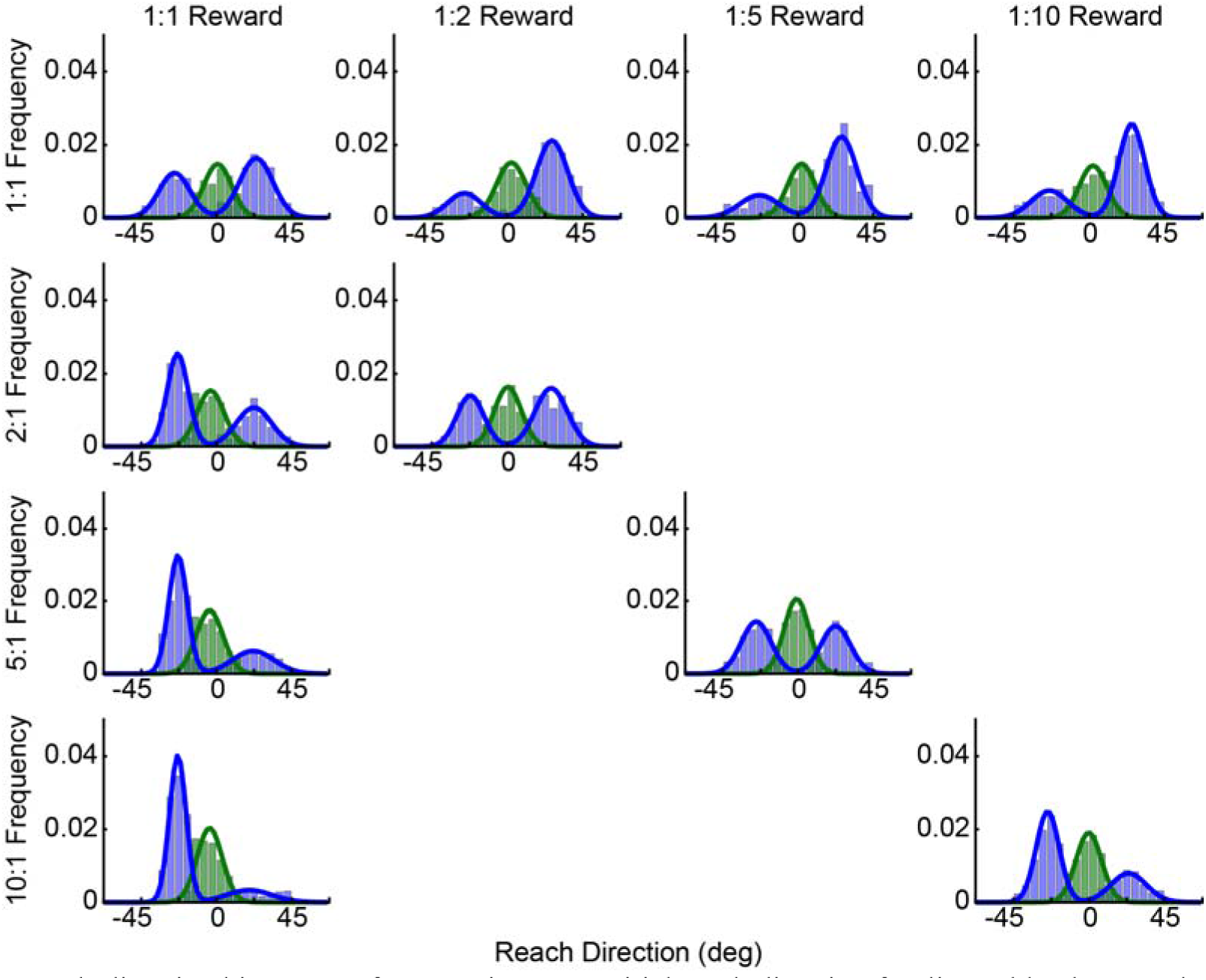
Reach-direction histograms for Experiment 1. Initial reach direction for direct (blue bars) and intermediate (green bars) reaches (data pooled across all participants), with varying reward and frequency ratios. In all cases, the data have been aligned such that reaches to the more rewarded target are in the positive direction (rightward), and reaches to the more frequent target are in the negative direction (leftward). As the relative reward ratio increases (left to right), reaches are biased toward the more rewarded target. Likewise as the relative frequency ratio increases (top to bottom), reaches are biased toward the more likely target. Fits reflect the average distribution parameters from bootstrapped fits to subsets of the pooled group distributions.

**Figure 3.**
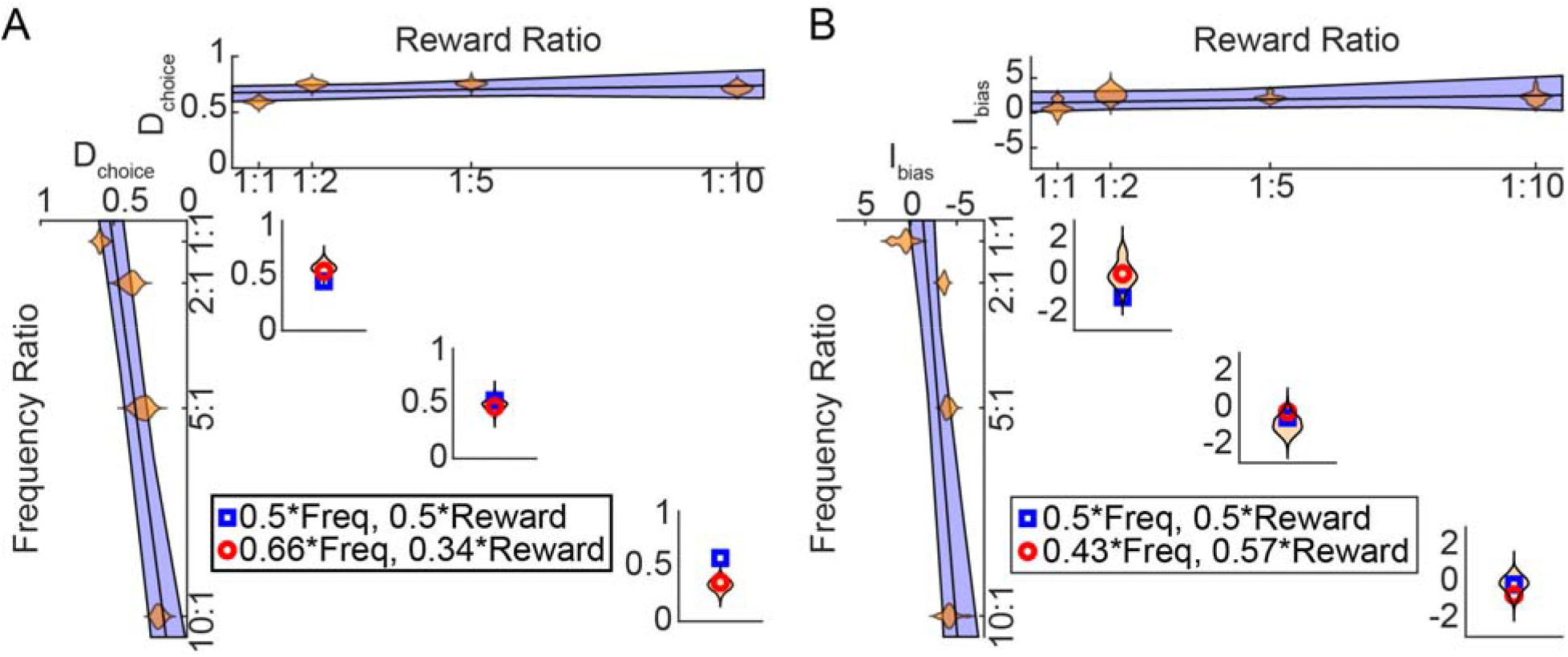
Estimation of biases in Experiment 1. For direct reaches (A) and intermediate reaches (B), the effect of changing reward (top) or frequency (left side) alone was fit by a linear regression. These regressions were then use to predict the reach direction bias observed when both reward and frequency were changing (3 small panels). Weighting the influence of frequency and reward unequally best explained the data (compare the red circles reflecting the best weighting of frequency and reward, to the blue squares reflecting equal weighting of frequency and reward). These weights indicated a much stronger influence of frequency compared to reward for direct reaches, but these factors were weighted differently for intermediate reaches.

Based on these effects of frequency and reward alone, we predicted how reaches would be biased when the effects of reward and frequency opposed each other. Specifically, the regressions above provided estimates of the relationship between frequency ratio and reach bias, and the relationship between reward ratio and reach bias. We then asked whether a (weighted) linear combination of these effects could explain the observed bias when frequency and reward both varied (see Methods). We noted that rather than weighing frequency and reward equally, we could explain the observed biases best when the relative influences (weightings) of reward and frequency were allowed to be unequal (where the reward weight equaled one minus the frequency weight; Fig. 3 A,B). For direct reaches, the best predictions were obtained if the influence of frequency on reach-direction bias was nearly twice that of reward (weight of frequency, 0.66 ± 0.11 SD; significantly different from 0.5, *t*(999) = 44.90, *p* = 8.21e-242). In contrast, for intermediate reaches we observed the best predictions when frequency was weighted slightly less than reward (weight of frequency, 0.43 ± 0.08 (SD); significantly different from 0.5, *t*(999) = −25.88, *p* = 1.93e-113). Overall, the weighting of frequency over reward was significantly greater for direct versus intermediate movements (Fig. 4A; weight difference = 0.23 ± 0.11 SD; paired t-test, *t*(999) = 64.47, *p* = 0.00; Kolmogorov-Smirnov test, *p* = 1.99e-272). These results agree with a second approach borrowed from neuroeconomics in which we assumed that the bias arose from a comparison of the relative subjective values of the two targets (see Supplemental Methods and Supplemental Figure 1). Such an analysis revealed that reward information was discounted less when estimating subjective value for intermediate reaches compared to direct reaches, consistent with its increased weighting for intermediate reaches as noted above.

**Figure 4.**
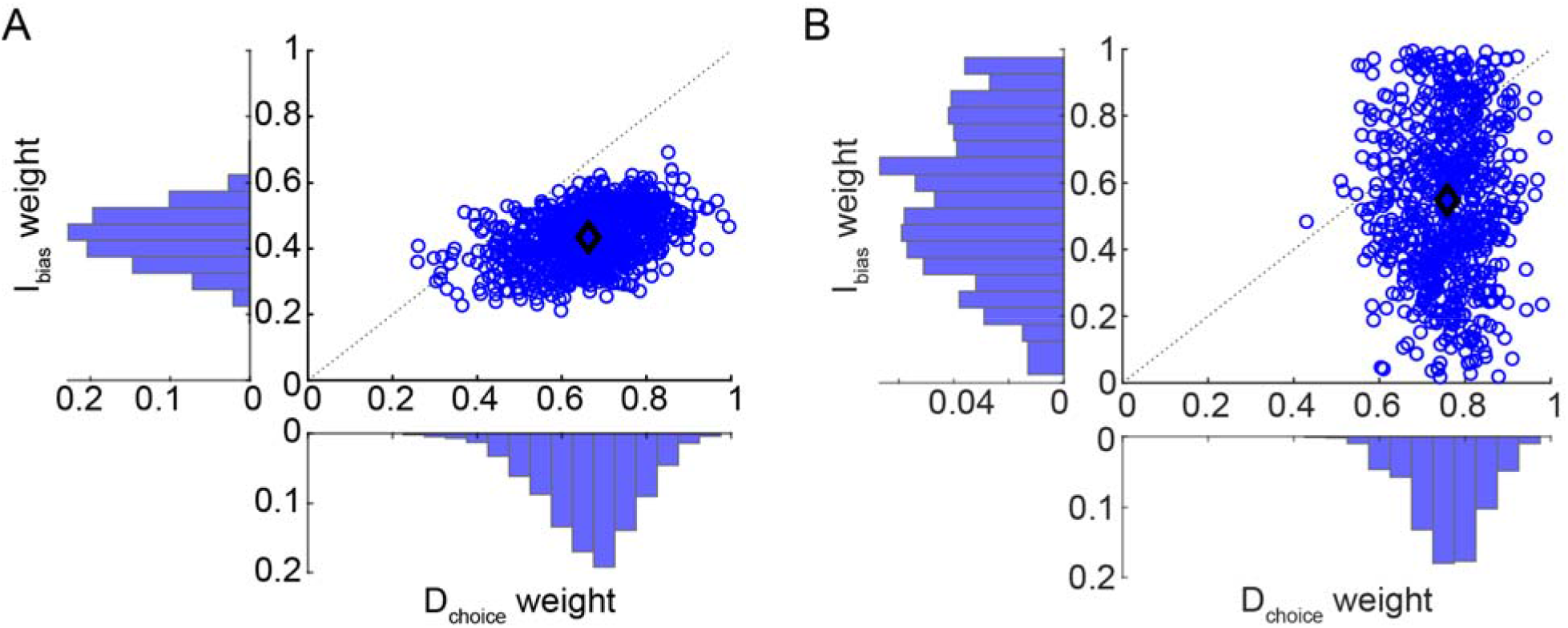
Comparison of likelihood-reward weight estimates for direct versus intermediate reaches, for Experiments 1 and 2. (A) The relative weighting of frequency for direct (D_choice_ weight) and intermediate (I_bias_ weight) reaches, and their relationship (blue circles), is plotted for Experiment 1. On average (black diamond), frequency tended to be weighed more heavily than reward for direct reaches, but the opposite weighting was observed for intermediate reaches. (B) The same weight distributions and relationships are plotted for Experiment 2. Although there was much greater variability of probability-reward weight estimates for intermediate reaches in this experiment, on average weighting effects were analogous to those observed in Experiment 1 (compare the black diamond in panels A and B).

### Prospective probability biases behavior analogously to frequency

One concern with modulating target frequency is that participants have to learn the relative frequencies between the two targets in order for that information to modulate behavior. Although the data suggest that varying the relative target frequencies did indeed change how people biased their responses, it is possible that participants did not notice the frequency manipulation and thus the biases observed were not actually representative of what happens when people consider the relative likelihood of the options in their decisions. In addition, developing a frequency bias meant that for a given individual there was no way to counterbalance the more frequent target within a block; thus there may have been idiosyncratic preferences, use-dependent repetition effects, or unintentional biomechanical biases unrelated to the frequency manipulation that influenced behavior. To address these concerns, we conducted a second experiment in which participants were explicitly shown the probability of the two targets alongside the reward values (Fig. 1B); each trial had no dependence on the previous one, and all reward and prospective-probability combinations were counterbalanced such that no target appeared more frequently within a block.

Despite these changes, on average participants exhibited analogous behavior to that of Experiment 1 (see Supplemental Fig. 2). As before, bootstrapping was used to examine the relationship between direct and intermediate movement biases arising from varying target reward and likelihood ratios. For direct reaches (Supplemental Fig. 3A), we again observed a bias in the proportion of reaches aimed to the more rewarding target due to the reward ratio (slope = 0.009 ± 0.021 SD) and a bias in the proportion of reaches aimed to the less probable target due to the probability ratio (slope = −0.11 ± 0.010 SD). For intermediate reaches (Supplemental Fig. 3B), we also observed a bias in the initial reach direction due to the reward ratio (slope = 0.093 ± 0.23 SD) and a bias in the reach direction due to the probability ratio (slope = −0.45 ± 0.39 SD).

In attempting to predict the relative influence of probability and reward, we observed that as with Experiment 1, direct reaches tended to be more strongly influenced by likelihood information compared to reward information (weight of probability, 0.76 ± 0.08 (SD); *t*(769) = 85.27, *p* = 0.00). Intermediate reaches were also biased by probability more than reward (weight of probability, 0.55 ± 0.24 (SD), *t*(769) = 5.68, *p* = 1.93e-08), although as before the influence of probability was significantly greater for direct reaches compared to intermediate reaches (Fig. 4B; paired t-test, *t*(769) = 23.67, *p* = 1.71e-93; Kolmogorov-Smirnov test, *p* = 3.3902e-101). These results closely resemble those observed in Experiment 1, replicating our prior findings that likelihood and reward information are weighed to different extents in direct and intermediate reaches. This suggests that direct and intermediate reaches reflect qualitatively different aspects of behavior.

### Risk/reward-seeking attitude modulates behavioral biases

In Experiments 1 and 2 (and historically in the literature), analyses were performed at the group level. This is largely because individual participants are idiosyncratic in their behavior: even in cases where the reward and likelihood of the two targets are balanced, some participants produce only intermediate reaches, some produce only direct reaches, and some produce a mixture of both. Such variable responses make it challenging to directly compare behavior in direct and intermediate reaches within a single subject. Moreover, it is not clear what may be driving an individual’s particular choice of movement strategy. To investigate these questions, we designed a new experiment that encouraged individuals to generate a mixture of direct and intermediate reaches in every condition (Fig. 5A). Specifically, we gave participants an explicit “intermediate” movement option on each trial by presenting an additional pair of targets farther from the start position, and modulated the relative rewards on offer between the direct (near) and intermediate (far) options (while keeping the left-to-right reward and frequency ratios fixed across the direct and intermediate options). This effectively created varying situations that could favor making direct or intermediate reaches (Fig 5A; see Methods), enabling the relative weighting of frequency and reward to be estimated for each individual (Fig. 6, 7). Visual inspection revealed that individuals exhibited similar trends as observed in Experiments 1 and 2: an increasing bias toward the more rewarded target as the reward ratio increased, and an increasing bias toward the more frequent target as the frequency ratio increased.

**Figure 5.**
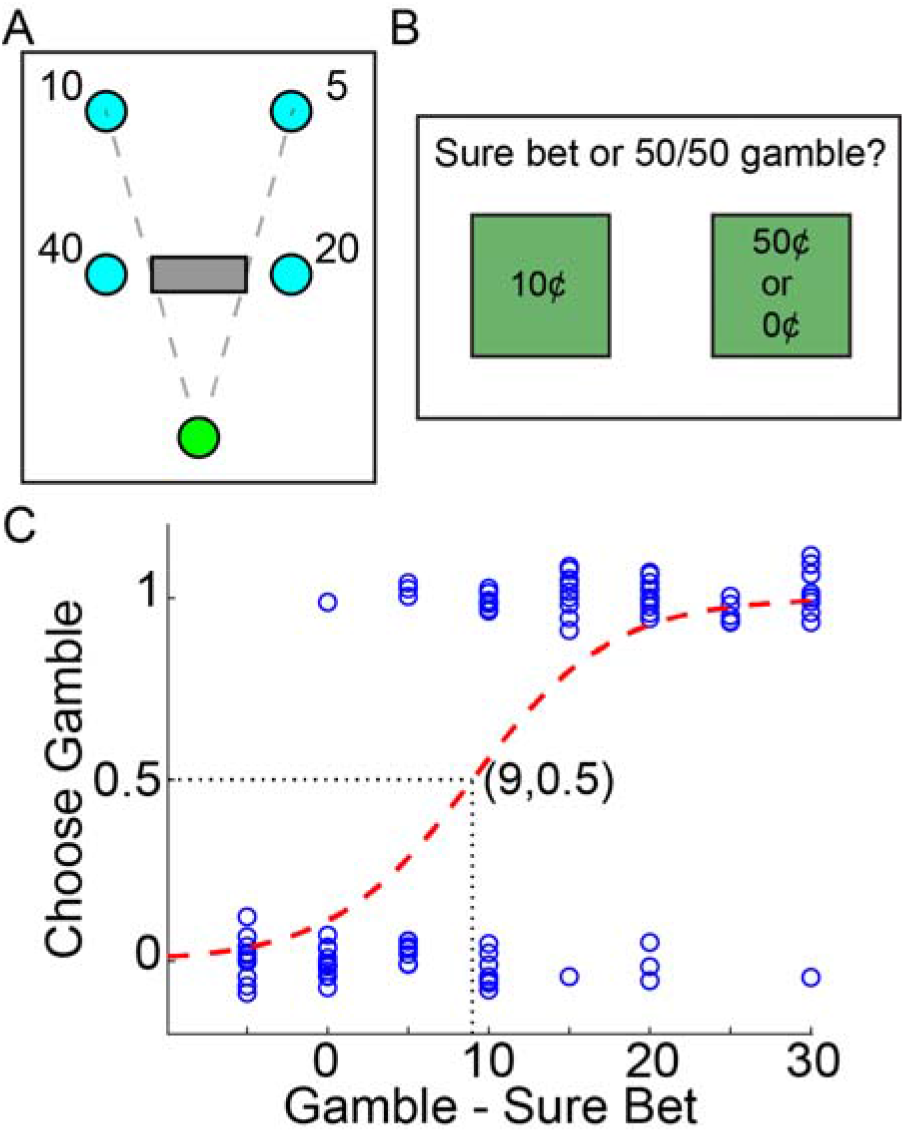
Methods for Experiment 3. (A) Participants completed a modified go-before-you-know task in which intermediate reaches were explicitly indicated. Specifically, starting from a home position (green), participants could reach directly to one of two near targets with no knowledge of which target was correct until after hitting the target, or they could pass through a rectangle that would cause both near targets and one of the far targets to disappear, revealing the correct far target. Both leftward or both rightward targets were “correct” on a given trial, and were assigned the same ratio of reward and likelihood. The reward offered for the far targets was a proportion of the reward available on the near targets; varying this proportion changed the relative desirability of making direct or intermediate reaches. The dashed lines were not visible to participants, but have been drawn to indicate the width of the “intermediate” rectangle relative to the far targets. (B) Participants also completed a utility task in which on each trial they indicated with a button press whether they preferred a sure bet or a 50/50 gamble. (C) Psychometric curves were fit to the choices made in the utility task, and the indifference point was estimated. Rightward shifts of the indifference point reflect greater risk aversion; leftward shifts reflect greater risk/reward-seeking tendency. Vertical jitter of data points was added for clarity.

**Figure 6.**
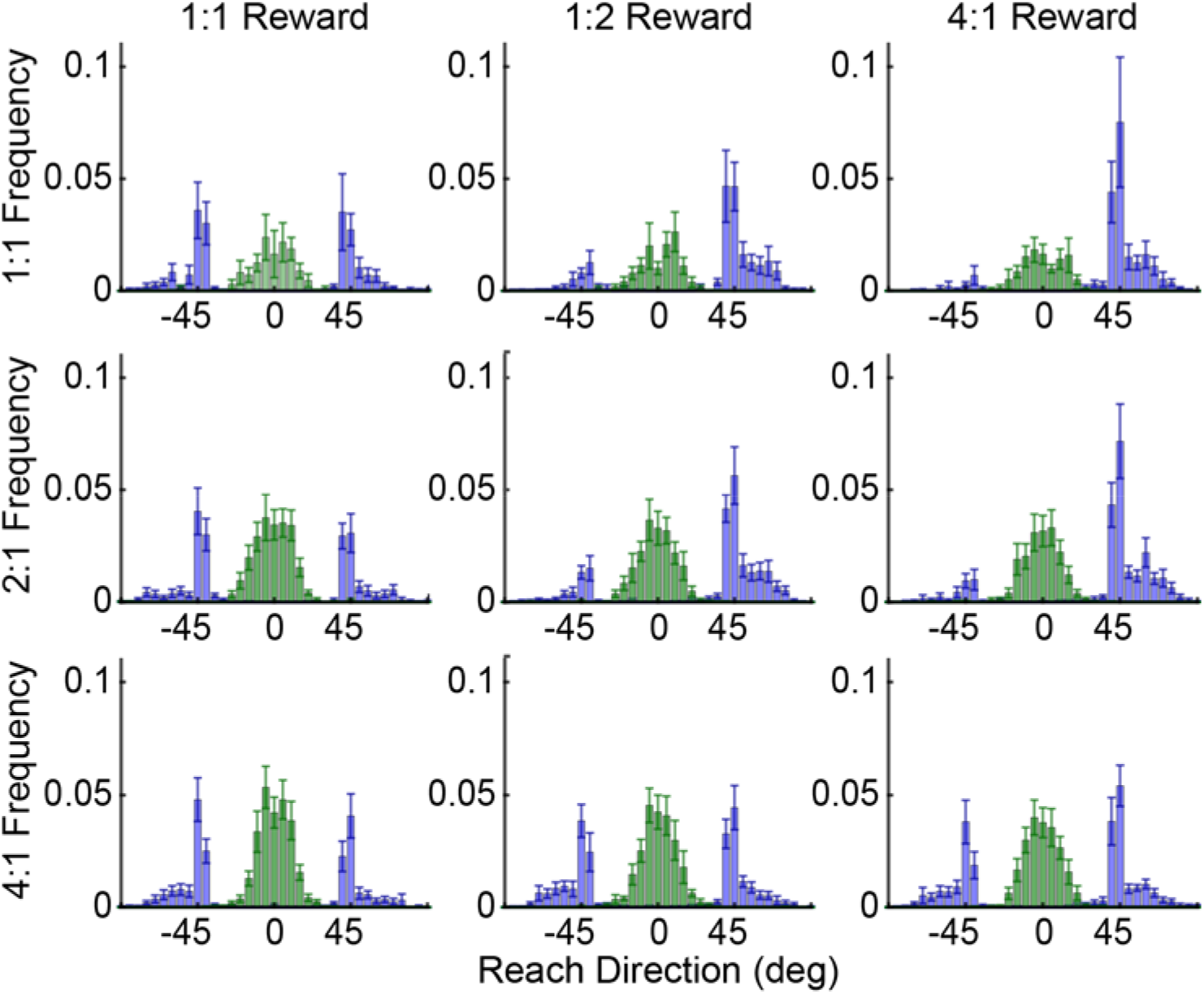
Average reach-direction histograms for Experiment 3. For each condition, individual histograms were generated for each participant and then averaged. Bar height reflects the proportion of total reaches aimed in a particular direction (bins of 5°) for a given individual in that condition; error bars reflect SEM across participants. Data were labeled according to whether individuals intercepted the rectangular “intermediate” target to hit the far targets (green bars), or reached to one of the near “direct” targets (blue bars). As in Fig. 3, data were sorted such that the positive reach directions were in the direction of the more rewarded target, while negative reach directions were in the direction of the more frequent target. As the relative reward ratio increased (left to right), reaches were biased toward the more rewarded target. Likewise as the relative frequency ratio increased (top to bottom), reaches were biased toward the more likely target. The skew in the direct-reach distributions likely arose because the intermediate “box” option was quite wide and participants had to avoid hitting it when making a direct reach.

**Figure 7.**
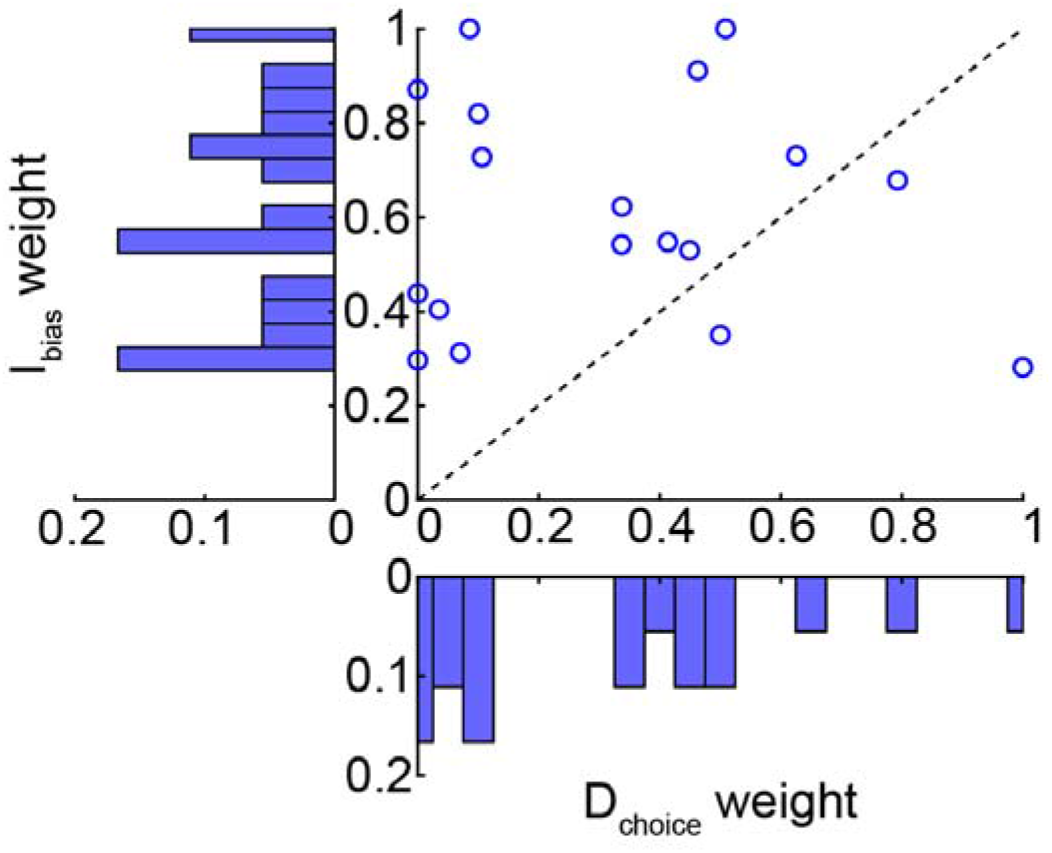
Comparison of individual likelihood-reward weight estimates for direct and intermediate reaches. As before, weights differed for direct and intermediate reaches, with no apparent systematic relationship evident.

For each participant, we estimated the relative influence of the frequency and reward ratios on reach-direction biases, for both direct and intermediate reaches (Fig. 7). In 3 participants, we were not able to fit a reasonable set of weights for both direct and intermediate reaches (i.e., it was likely that their behavior was idiosyncratic in one or more conditions). Across the remaining participants, frequency and reward influenced behavior to an unequal extent for direct reaches (weight of frequency, 0.32 ± 0.30 SD; *t*(17) = −2.52, *p* = 0.02), but not for intermediate reaches (weight of frequency, 0.62 ± 0.24 SD; *t*(17) = 2.02, *p* = 0.06). More importantly, the relative influence (weight) of frequency and reward for direct and intermediate reaches was significantly different from each other (paired t-test, *t*(17) = −3.16, p = 0.006). However, the relationship between these weights was not systematic across individuals (no correlation between the weights for direct and intermediate reaches, *r*^2^ = 0.002, p = 0.85). This lack of relationship between the weights estimated for direct and intermediate reaches was unlikely to be an artifact of the analysis method, as suggested by a parameter recovery analysis (see Supplemental Methods and Supplemental Fig. 4).

Next, each individual’s risk/reward-seeking tendency was assessed to examine if that could explain observed individual differences in behavior (Fig. 5B,C). We noted one outlier who exhibited unusually strong risk-seeking behavior; to ensure that this person did not skew our findings we removed this individual from further analysis. We then examined whether risk/reward-seeking tendency might influence the degree to which individuals generated direct or intermediate reaches (Fig. 8A). We observed a significant correlation between the proportion of intermediate movements generated on average across conditions and the tendency to be risk/reward-seeking (r^2^ = 0.47, p = 0.011): risk/reward-seeking individuals tended to produce more intermediate movements. Interestingly, the risk-seeking outlier falls very close to the estimated regression line (r^2^ = 0.50, p = 0.005). This result, taken alongside the findings from Experiments 1 and 2 showing that intermediate movements were more strongly biased by reward compared to direct movements, suggest that intermediate movements reflected an attempt to maximize rewards earned during the task. Consistent with this idea, on average across all task conditions, individuals who were more risk/reward-seeking also tended to modulate their behavior to a greater extent (i.e., exhibited a greater shift from making intermediate to direct reaches) as the near targets became relatively more rewarding compared to the far targets (Fig 8B; *r*^2^ = 0.27, p = 0.034). Results from all three experiments together suggest that intermediate and direct movements seem to reflect attempts to maximize reward versus success respectively, and this behavior seems to arise from the degree to which an individual is risk/reward-seeking.

**Figure 8.**
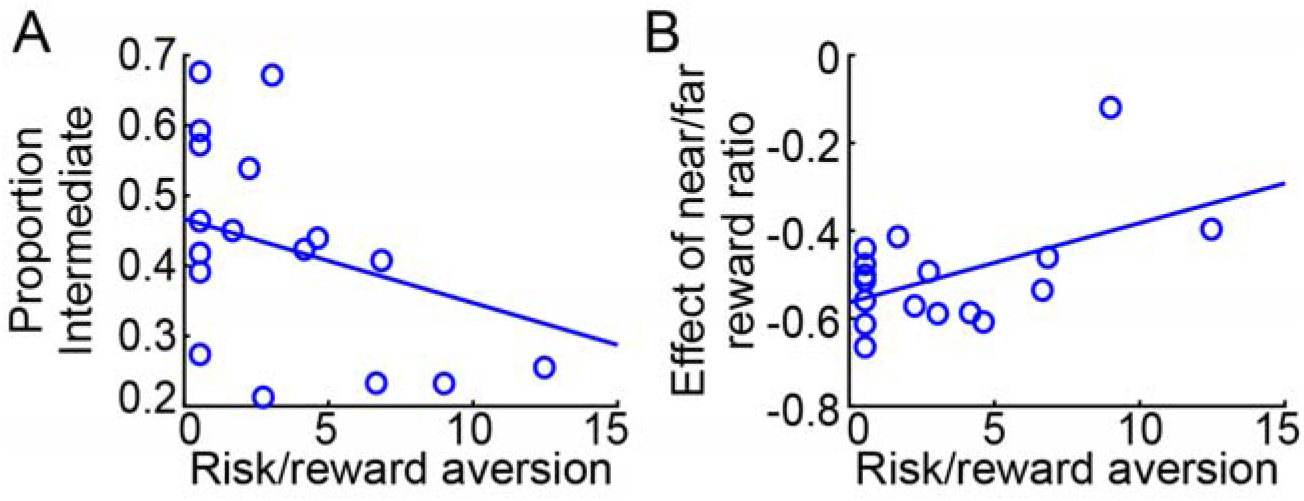
Risk-seeking tendency explained individual differences in behavior. (A) The average proportion of intermediate movements generated correlated with a tendency to be risk/reward seeking, suggesting intermediate movements reflect a strategy to maximize reward. (B) Risk/reward-seeking tendency increased the degree to which the relative reward ratio of near to far targets shifted the proportion of intermediate reaches produced.

## DISCUSSION

This study examined two key questions regarding how people respond when faced with multiple potential options. The first question examined how individuals weigh reward and likelihood information (i.e., exogenous and endogenous rewards) in determining the relative desirability of each option, and whether any preference biases that arose would be analogous for both direct and intermediate reaches. We showed that the decision biases induced by varying the desirability of the two targets were not analogous for direct and intermediate reaches. Specifically, direct reaches were primarily influenced by target likelihood, while intermediate reaches were more influenced by reward. This importantly suggests that intermediate reaches do not arise from the averaging of direct-reach plans. Instead, these findings are consistent with and extend previous work demonstrating that intermediate movements are deliberately adopted as a distinct motor strategy to improve task performance (Alhussein and Smith, 2021; Dekleva et al., 2018; Wong and Haith, 2017) – in this case, to maximize rewards. More broadly, these findings reveal that different behaviors, even when generated within the context of the same task, can be differentially motivated by reward or success; or equivalently, that exogenous rewards (money) and endogenous rewards (success) can influence behavior differently. This highlights the fact that not only are reward and likelihood generally subjectively viewed as differentially important in determining a movement response (see Supplemental Fig. 1; Kahneman and Tversky, 1979; Onagawa et al., 2019), but moreover that their subjective importance changes depending on the strategy with which one chooses to approach a given task.

Direct and intermediate movement strategies therefore appear to be optimizing different things: task success versus reward outcomes, respectively. Such an optimization makes sense: on direct reaches, individuals must commit to a decision before movement onset; hence the best approach is to choose the target that is more likely to be correct, even at the cost of occasionally missing large, rare rewards. Direct reaches reflect a relatively low-effort, low-risk strategy to successfully hit the correct target most of the time. On the other hand, intermediate reaches allow individuals to delay commitment to a particular option until more information can be acquired, and affords the ability to change one’s mind. Because this strategy reduces the risk of failure, individuals can instead focus on maximizing rewards; hence it is sensible to more equally weigh the likelihoods and the rewards of the two options. However, intermediate movements are also more effortful to complete: they necessarily require constant monitoring of the target options throughout the movement to identify the correct target as soon as it is revealed, and they always require a motor correction to be generated midway through the movement. This taxes both cognitive and physical resources. It has previously been shown that the prospect of larger rewards can offset the cost of physical and mental effort (Dixon and Christoff, 2012; Manohar et al., 2015; Schmidt et al., 2012; Summerside et al., 2018). By improving reward outcomes, an intermediate-movement strategy can thereby offset the additional cost of generating such movements. Thus, direct and intermediate reaches appear to reflect distinct movement strategies to respond to decision uncertainty, residing on opposite ends of the effort-reward (and risk-reward) trade-off.

Our second question examined whether we could explain why individuals preferentially generate direct and intermediate movements to different extents when asked to reach under the same conditions of target uncertainty. Given that direct and intermediate reaches seem to be distinct strategies favoring success or reward respectively, we hypothesized that individual risk/reward-seeking tendencies might influence how individuals choose to respond (Wong and Haith, 2017). At first glance, one might think that because intermediate movements enable individuals to change their mind, this might be the strategy preferred by more risk-averse individuals. Instead we found the opposite result: the more risk/reward-seeking the individual, the greater the proportion of intermediate reaches they generated. Since intermediate movements were more strongly influenced by reward relative to direct reaches, however, this makes sense: reward-seeking individuals preferentially adopt a strategy that allows them to maximize their potential rewards.

Note that even though intermediate movements have the potential to also improve success at hitting the correct target, not every intermediate movements is successful (see for example, Wong and Haith, 2017); thus intermediate movements are not necessarily the more risk-averse movement strategy. Additionally, as noted above, they are also the more effortful strategy, and thus by subjectively valuing rewards to a greater extent individuals may be more likely to perceive that they are achieving an adequate offset for this additional effort cost (Galaro et al., 2019; Klein-Flügge et al., 2016). Consistent with these ideas, risk/reward-seeking individuals also responded more strongly to the reward manipulation in Experiment 3 that was designed to modulate the production of intermediate versus direct reaches (i.e., the relative rewards offered for near versus far targets). In contrast, individuals who were more risk-averse and less reward-seeking seemed to favor making direct reaches, suggestive of an effort to maximize success at hitting the correct target.

Interestingly, our results imply a hierarchical decision process in which, in response to choice uncertainty, individuals first select a movement strategy (i.e., whether to generate a direct or intermediate reach). This movement strategy in turn may then modulate the way in which reward and likelihood information bias reach choice. Such a hierarchical decision structure has previously been argued to be favorable for action selection (Rosenblatt and Payton, 1989; Tyrrell, 1993). Interestingly, such a hierarchy results in two consequences that are relevant here. First, it allows relevant sensory information to be introduced at lower levels of the hierarchy after some decisions have already been made, avoiding a sensory bottleneck and maintaining a greater amount of information in the network. Second, it enables individuals to identify and select “compromise candidates”, or options that are favorable given overriding constraints. In our case, the compromise candidate is reflected in the intrinsic movement bias arising after a decision strategy has been adopted; moreover, subject-specific movement-strategy preferences can be determined according to an individual’s risk/reward-seeking propensity without first needing to consider the reward and likelihood values on offer in a given trial. Indeed, we note that while likelihood and reward information seem most relevant for setting movement biases once a choice of direct or intermediate has been made, the propensity for risk/reward-seeking modulates decisions at multiple levels of the hierarchy: affecting the preference for making direct versus intermediate reaches, and subsequently influencing the relative weighting of likelihood and reward information in determining which option is more desirable. Importantly, such a hierarchical structure suggests that the subjective value placed on action choices – in neuroeconomics terms, their utility – not only influences movement decisions about where to reach (Wolpert and Landy, 2012), but is itself influenced in turn by movement decisions about how to reach.

While some amount of effort has been made to investigate the neural underpinnings of direct and intermediate movements (Cisek, 2007; Cisek and Kalaska, 2002; Dekleva et al., 2018), much of this work thus far has focused on premotor cortex. In contrast, efforts to study the representation of subjective value and relative choice desirability have largely investigated midbrain structures including the basal ganglia and associated areas of frontal cortex (Croxson et al., 2009; Doi et al., 2020; Fox and Poldrack, 2009; Klein-Flügge et al., 2016; Levy et al., 2010; Samejima et al., 2005; Schmidt et al., 2012; Shin et al., 2021; Trepel et al., 2005). Intriguingly, however, there is some suggestion that parietal reach region in the posterior parietal cortex might be a potential area where information about motor goals and goal desirability converge (Trepel et al., 2005). Specifically, neurons in parietal reach region have been shown to modulate according to both the rewards and likelihoods of action choices as if computing target utility (Platt and Glimcher, 1999); additionally, this region seems capable of simultaneously representing multiple potential movement goals (Klaes et al., 2011). Subjective utility has also been shown to modulate cortical excitability in M1 and directly influence behavior (Galaro et al., 2019). This suggests multiple points at which information about target desirability can interact with decisions about where to reach, in line with the notion of a hierarchical decision structure (Rosenblatt and Payton, 1989; Tyrrell, 1993).

In conclusion, our data suggest that when faced with moving in the face of uncertainty, individuals deliberately adopt a motor strategy in line with their proclivity to be risk/reward-seeking. In particular, this proclivity influences whether individuals prefer to commit to a guess before moving (i.e., generating direct movements), versus hedging one’s bets and delaying commitment until later in the movement (i.e., generating intermediate movements). Such findings support the idea that intermediate movements are planned as part of a deliberate movement strategy to maximize exogenous reward, distinct from that underlying the planning of direct reaches which seeks to maximize endogenous reward (i.e., success). Moreover, the choice of movement strategy appears to influence the manner in which individuals evaluate the subjective value of the available options. Together, this suggests that in the face of uncertainty, people trade off maximizing task success (endogenous rewards) and earning monetary rewards (exogenous rewards) depending on their individual risk/reward attitudes, and these decisions influence both the choice of movement strategy as well as the apparent subjective value of the available options.

## METHODS

Sixty-nine right handed adult neurotypical individuals were recruited for this study. Of those individuals, 24 participated in Experiment 1 (age 18-40, average 26.0 ± 5.1; 17 female), 17 participated in Experiment 2 (age 22-34, average 24.1 ± 2.8; 16 female), and 28 participated in Experiment 3 (age 21-33, average 24.8 ± 2.4 years; 22 female). One participant in Experiment 3 was unable to complete the study due to scheduling constraints, so data from only 27 participants was available for analysis. Of these 27 individuals, 18 participants reliably generated both direct and intermediate reaches in every condition (i.e., across the varying reward and frequency ratios) such that the relative weighting of frequency and reward could be estimated; the analyses here thus reflect the data from these 18 individuals. Sample sizes were chosen to be comparable to or greater than previous studies utilizing a similar task (e.g., Chapman et al., 2010; Wong and Haith, 2017). All participants were naïve to the purposes of this study, and provided written informed consent. Experimental methods were approved by the Einstein Healthcare Network Institutional Review Board and participants were compensated for their participation as a fixed hourly payment ($12/hr) plus the potential to earn an additional bonus (up to $6/hr) based on performance in the task.

Participants made planar reaching movements while seated in a Kinarm Exoskeleton Lab (Kinarm, Kingston, ON, Canada). Hands, forearms, and upper arms were supported by the robot in troughs appropriately sized to each participant’s arms, and the linkage lengths of the robot were adjusted to match limb segment lengths for each participant to allow smooth motion at the elbow and shoulder. Vision of the arm was obstructed by a horizontal mirror, through which participants viewed a cursor (1 cm diameter) representing the position of the index finger and targets (2 cm diameter) displayed by an LCD monitor (60 Hz) in a veridical horizontal plane. Movement of the arm was recorded at 1000 Hz.

### Experiment 1

Participants were asked to complete a “go before you know” task (Chapman et al., 2010; Wong and Haith, 2017) in which after a brief random delay (900 ms ± 600 ms) two potential targets (rings, 2 cm diameter) were presented 45 deg apart, 20 cm from a central starting position at the beginning of each trial (Fig. 1A). Participants were free to move any time after the targets appeared, and had 2500 ms from the time of target appearance to complete their reach by shooting through the target. Once participants began their movement (moved 4 cm from the starting position or exceeded a hand velocity of 0.1 m/s), the correct target (i.e., the one that had to be hit to earn a reward) was revealed by becoming solid in color and the distractor target disappeared. Participants were rewarded for hitting the correct target while maintaining a minimum hand velocity (0.3 m/s) without exceeding the vertical target distance from the starting position before hitting the target. Successful target interception resulted in a reward, the amount of which was determined according to an initial Utility Assay (see below for details). If participants moved too slowly, a “Too Slow” message was displayed at the end of the trial and participants did not earn a reward for that trial.

Each target was associated with a reward and a likelihood, which varied per block. Monetary rewards were set as ratios of 1:1, 2:1, 5:1, and 10:1, and the reward amount assigned to each target in cents was displayed to the participant on every trial (Fig. 1A). Each target was also associated with a likelihood according to the relative frequency with which that target was correct in each block; frequency ratios were 1:1, 2:1, 5:1, and 10:1. Participants were not informed that the targets would vary in frequency, but could learn these frequencies through experience. In all cases, the more rewarding target was always the less frequent target. Therefore within a single block, the more rewarding target always appeared on the same side; the more-rewarding (lower-frequency) target side was counterbalanced across blocks and participants. Since target frequencies were learned by experience, participants always completed all blocks in the same order: increasing reward ratios at a frequency ratio of 1:1, followed by increasing reward ratios at a frequency of 2:1, etc. Because target frequencies varied between blocks, blocks consisted of 30-44 trials each. At the end of the session, participants were paid a fraction of the total rewards earned throughout these blocks, and thus were encouraged to treat all trials as if they would actually earn the reward offered for hitting the correct target.

Prior to this session, participants completed a Utility Test (Fig. 5B). Since not all people value fixed monetary amounts to the same extent, in Experiments 1 and 2 an effort was made to equate reward utility across participants such that all individuals would be similarly motivated by the individual target rewards. To do so, participants completed a pretest in which on every trial they were presented with two options, a fixed monetary amount (“Sure Bet”, e.g., 10 cents) and a 50-50 gamble (“Gamble”, e.g., $1.00 or 0 cents). Participants had to choose which they would prefer; to have the fixed amount of money or take a gamble and win either of the two amounts depending on the outcome of a fair coin flip. Participants were not informed of the outcomes of their choices until the end of the block, where they saw a total score reflecting the sum of all their choices. Hence, success or failure of the gamble option did not impact the decision on the next trial. Participants received a fraction of the total earned as part of their final monetary compensation; thus they were instructed to treat each trial as if they would actually earn the outcome of that trial. Immediately following this block, we estimated the likelihood of choosing a fixed amount versus the expected outcome of the gamble for each presented pair of options, and fit a plane to the data; this plane was used to determine the monetary value for which participants were equally likely to choose that value versus a gamble with an expected outcome of 10 cents (i.e., the subjective point of equivalence). If the estimated value was less than 5 cents or greater than 20 cents we adjusted the value to be within this range to avoid offering excessively large or small reward values to participants. This monetary value was then set to be the base reward value in the go-before-you-know experiment above from which all the reward offers were computed (as multiples of the base reward value). Code to run this experiment is available on our lab Github page (https://github.com/CML-lab/KINARM_GBYK_RwdFreq).

### Experiment 2

Experiment 2 was identical to Experiment 1 except that instead of varying the frequency associated with each target, we varied the target probability on a trial-by-trial basis (such that there was an equal long-run likelihood of the right or left target being correct and the blockwise expected value of the two targets was equal, but on a particular trial each target was explicitly associated with a particular probability of being correct). Participants were informed of these probability ratios based on the appearance of the target; targets consisted of 2-cm squares divided into a checkerboard pattern, where the ratio of randomly chosen red to blue pixels in the target represented the target probability of being correct (Fig. 1B). Target probabilities could be 50:50, 70:30, or 80:20; twenty trials at each probability ratio were randomly intermixed within a single block for a total of 60 trials each. Reward ratios could be either 1:1, 2:1, or 5:1; each block contained a single reward ratio for every trial, with all participants experiencing reward ratios that increased throughout the experiment to minimize any potential biases due to moving from high-reward-ratio to low-reward-ratio trials. To ensure participants had sufficient time to observe the stimuli (probabilities and rewards available), the targets and reward values appeared 900 ± 300 ms prior to a go tone indicating that the reaching movement could begin. As above, a Utility Test was performed and the result was used to set the base reward value on an individual-subject basis. Code to run this experiment is available on our lab Github page (https://github.com/CML-lab/KINARM_GBYK_RwdProb).

### Experiment 3

Unlike the first two experiments, Experiment 3 presented individuals with 5 targets at the start of each trial: a near pair of circular targets 90° apart and 10 cm from the start position (7.07 cm in depth), a rectangular box spanning 46° across the vertical midline and 7.07 cm from the start position in depth (i.e., in line horizontally with the near targets), and a far pair of targets located 45° apart and 15 cm from the start position (13.86 cm in depth; Fig. 5A). Thus in this Experiment, participants were presented with 2 explicit choices; they could either directly reach for one of the two targets closest to the start position, or they could pass through the rectangle on their way to the far targets(representing an explicit “intermediate” option). If participants reached toward one of the near targets, the targets remained on screen unchanged until one was hit, after which individuals received feedback on whether they hit the correct or incorrect target. In contrast, when participants passed through the rectangle, the closest pair of targets vanished as well as the incorrect far target, leaving only the correct far target visible for participants to steer their reach toward. To give participants time to assess the greater complexity of the visual display, participants were required to view the screen for at least 2 seconds before a tone sounded to indicate participants could begin moving.

The two targets on a given side of the midline were yoked together, such that the “correct” target was either the two targets on the right or the two targets on the left. That is, participants primarily had to choose whether to aim rightward or leftward. Participants were informed of this relationship between the targets. As in Experiment 1, correct targets appeared with differing frequencies in each block (with fixed frequency ratios of 1:1, 2:1, and 4:1); the side of the more frequent target differed across blocks within an individual, and was counterbalanced across participants. In addition, the reward ratio of the two near targets was the same as the reward ratio of the two far targets, in ratios of 1:1, 2:1, and 4:1. However, the pair of targets closer to the start position were worth anywhere from 1 to 4 as much as the two targets farther from the start position. Thus when the near targets were worth four times the value of the near targets, a direct reach strategy was implicitly encouraged; when the near targets were worth the same value as the near targets, an intermediate reach strategy was implicitly encouraged. The reward values of the four targets were explicitly written on the screen. Reward values (in cents) were fixed for all participants, with the more rewarding side counterbalanced across blocks and participants such that individuals could not predict from one block to the next which side would offer more reward or be more frequently correct. The overall result of this experimental design was to encourage individuals to generate both intermediate and direct reaches regardless of the exact target reward and frequency, moreover, we were able to explicitly identify reaches that were “direct” versus “intermediate” without resorting to model fitting or assumptions in the analysis (see below). A Utility Test was conducted (Fig. 5B), but the outcome of this test was not used to set the reward values during the session; instead, this test was used to estimate risk/reward sensitivity (see below). Code to run this experiment is available on our lab Github page (https://github.com/CML-lab/KINARM_GBYK_DiscreteChoice).

### Data Analysis

Reaches were analyzed offline using programs written in MATLAB (The MathWorks, Natick, MA, USA). Data and analysis code are available at https://osf.io/jn79v/?view_only=c6ae57430ce34bf99b0f3cd6ab213934. Movement onset was identified according to a velocity criterion (tangential velocity greater than 0.05 m/s in Experiments 1 and 2, 0.02 m/s in Experiment 3), and verified by visual inspection. The initial reach direction was determined as the direction of the velocity vector 100 ms after movement initiation. In Experiment 1, reaches were excluded from further analysis if the initial reach direction was greater than 45° away from the midline or the reach was not completed in 1000 ms from target appearance; this led to a median of 6.8% reaches being removed for each participant. In Experiment 2, reaches were excluded if the initial reach direction was greater than 45° away from the midline or the reach was not completed in 650 ms after the go tone; this led to a median of 11.4% of reaches removed. For Experiment 3, reaches were excluded if they were greater than 75° away from the midline, the reach was not completed in 2500 ms, or if the trial was incomplete; this led to a median of 10.5% of reaches removed in Experiment 3.

In Experiments 1 and 2, reaches were classified as “direct” or “intermediate” according to a time criterion. At each time point during the reach, we calculated for each target the instantaneous target direction as the direction in which the hand would need to move, given its current position, to be heading directly toward the target (see Fig. 1C). We also computed the actual reach direction of the hand at each time point according to the direction of the instantaneous velocity vector at that time. We identified the time at which individuals first committed to reaching toward one of the two targets (T_commit_) as the time when the reach direction was equal to the instantaneous target direction. We noted whether T_commit_ occurred before or after the time that the true target was revealed (plus an additional 60 ms to account for delays between when the command was sent to modify the display and when individuals might actually perceive the display changing; T_reveal_). If T_commit_ occurred prior to T_reveal_, we considered this to be a “direct” reach; otherwise the reach was classified as “intermediate”. Note that due to trial-to-trial variability in instantaneous hand velocity during the reach, this classification approach had no relationship to when the initial reach direction was determined, which was always measured a fixed 100 ms into the reach.

For Experiment 3, reaches were explicitly identified as direct or intermediate depending on which target the reach first intercepted. Whenever a reach intercepted the rectangle it was classified as intermediate, otherwise it was considered a direct reach. Due to the way the task was programmed, it was not possible for individuals to intercept both the rectangle and one of the near targets; hitting the rectangle caused the near targets to disappear leaving only the far targets as viable options, while hitting one of the near targets (or missing both near targets and the rectangle) caused the trial to end.

Classification of reaches (whether by a temporal criterion or direct examination) resulted in two reach-direction distributions: a bimodal distribution of reaches aimed directly to the two targets, and a unimodal distribution of reaches aimed intermediate between the two targets. For Experiments 1 and 2, due to idiosyncratic behavior across individuals, reach directions across all individuals were pooled prior to analysis. Reach-direction distributions were analyzed separately by fitting a bimodal or unimodal Gaussian using the built-in fitgmdist and the fitdist functions in Matlab respectively. Variance associated with the fit parameters was estimated using a bootstrap approach in which Gaussians were fit to a randomly selected subset of 75% of the participants, and this subsampling was repeated 1000 times to estimate distributions for each of the fit parameters. Iterations were removed when one or more fits did not return valid parameters, or when by random chance insufficient numbers of direct or intermediate trials were available to fit. Of particular interest were the ratio of the two modes of the bimodal distribution (*D_choice_*), which reflects the relative frequency with which individuals chose the higher-rewarded but less-likely target on direct reaches, and the shift of the mean of the unimodal distribution away from zero (*I_bias_*), which reflects a bias in the initial reach direction toward the higher-rewarded but less-likely target for intermediate reaches.

Separately for direct and intermediate reaches, the effect of reward and likelihood on reach-direction biases were estimated by fitting linear regressions to the parameter estimates (*D_choice_* or *I_bias_* respectively) when the likelihood or reward ratio of the two targets was 1:1 respectively. These regressions estimate how reward or likelihood independently modulated reach direction. These regressions were used to estimate the expected *D_choice_* or *I_bias_* values for the cases when both reward and likelihood were different between the two targets. In general, the *D_choice_* or *I_bias_* values calculated from an equal weighting (average) of reward and likelihood effects were not observed to fit the actual data well (based on visual inspection). Thus to identify the best relative weighting of reward and likelihood information that could explain the behavioral data, we searched for the weighted sum of independent reward and likelihood effects that best predicted the bootstrapped *D_choice_* or *I_bias_* values for each bootstrap iteration. These weights represented the relative contributions of reward and likelihood information influencing reach direction, for direct or intermediate reaches respectively. In all cases, values are reported as mean ± standard deviation (SD), unless otherwise noted to be reporting the standard error (SEM). Weight distributions were compared to 0.5 (equal weighting) using t-tests, and to each other using a Kolmogorov-Smirnov test.

For Experiment 3, because the task was designed to encourage both direct and intermediate reaches regardless of the combination of reward and likelihood, data were analyzed at the individual-participant level in a within-subjects analysis. For direct reaches, we computed *D_choice_* as the proportion of reaches aimed at the more rewarding target over the total number of successfully completed reaches. For intermediate reaches, *I_bias_* was estimated as the mean reach direction on intermediate reaches. As in Experiments 1 and 2, we used regressions to estimate the influence of reward or likelihood alone on *D_choice_* or *I_bias_* when only one factor was changing; then we computed the weighted sum of the influence of reward and likelihood that best explained the data in the cases when reward and likelihood for the two targets were different. If reward and likelihood exerted the same influence on direct and intermediate reaches, we would expect the relative reward and likelihood weights to be similar for these two types of reaches. We compared the similarity of these weights for direct and intermediate reaches using a paired t test and examined their correlation using linear regression.

The proportion of intermediate reaches generated by each individual was calculated by counting the fraction of intermediate reaches produced in each condition, then taking the average. The degree to which the relative reward values of the near to far targets modulated the proportion of intermediate reaches was measured by finding, for a given ratio of near-target reward to far-target reward within each condition, the difference in the number of intermediate versus direct reaches as a proportion of the total number of reaches. Then a regression was fit to examine the relationship between the near-to-far reward ratio and the difference in intermediate to direct reaches across all conditions. The slope of this regression was taken as an estimate of the degree to which the near-to-far reward ratio modulated the fraction of intermediate reaches; steeper slopes (more negative values) indicated individuals who produced more direct compared to intermediate reaches when the near targets were more rewarding than the far targets, compared to when the near and far targets were equally rewarding.

Risk/reward sensitivity was also examined in Experiment 3 using data from the Utility Test. For each participant, the choice (coded as 1 if choosing the Gamble and 0 if choosing the Sure Bet) was plotted against the reward difference (the expected value of the gamble minus the value of the sure bet) on that trial (Fig. 5C). Then, a psychometric curve was fit to the data, and the indifference point was estimated as the point where the curve crossed 0.5 (i.e., an equal chance of choosing the Gamble versus the Sure Bet). Larger indifference points reflect a more risk-averse (requiring a larger difference between the expected value of the Gamble and the value of the Sure Bet in order to choose the Gamble) and less reward-seeking (favoring the Sure Bet despite the greater potential reward to be earned by choosing the Gamble) attitude.

## ACKNOWLEDGEMENTS

The authors thank Adrian Haith and Samuel McDougle for helpful discussions.

