## Supplemental Materials for "Motor plans under uncertainty reflect a trade-off between maximizing reward and success"

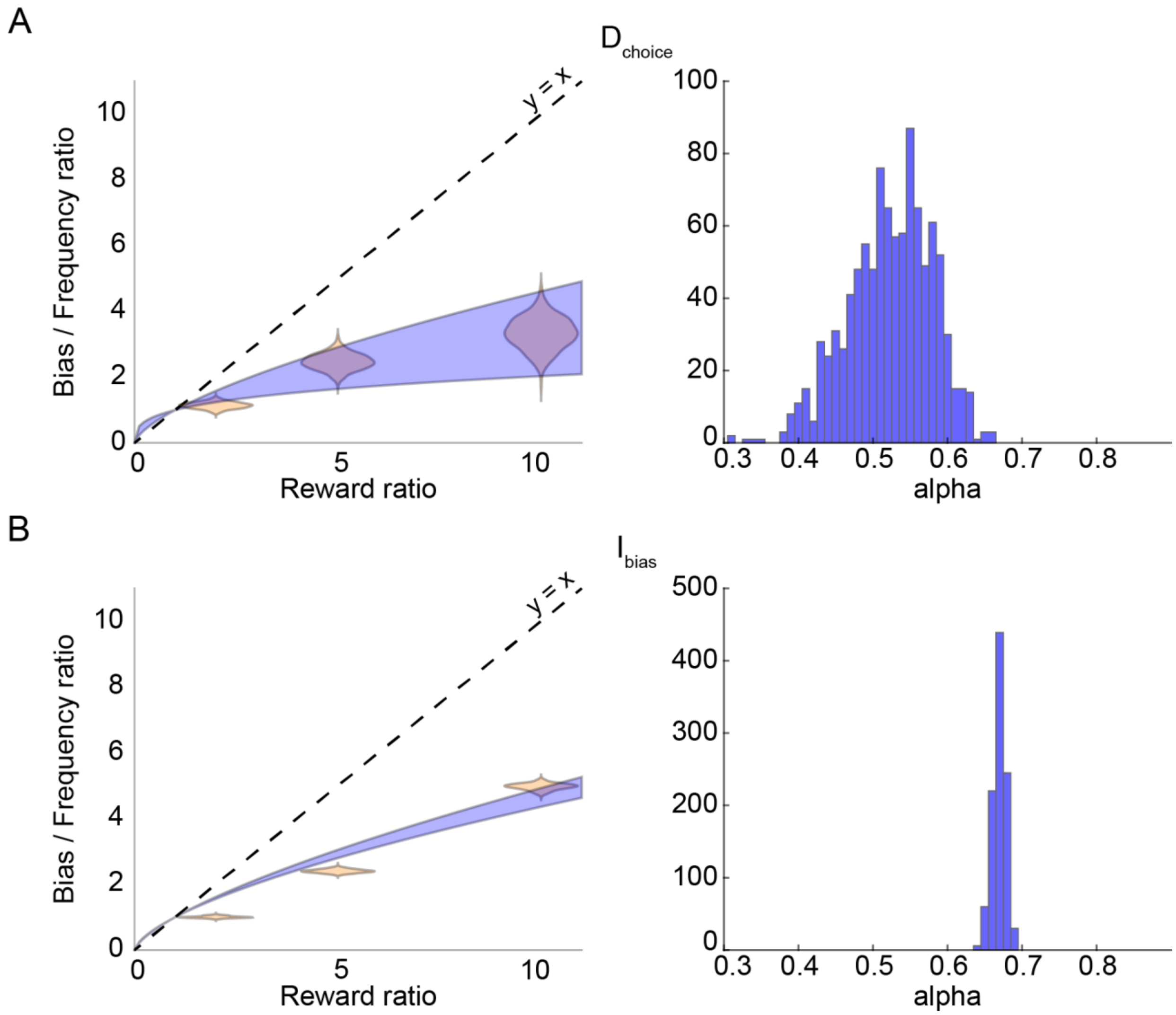


**Supplemental Figure 1.** Subjective valuation fits for Experiment 1. In addition to estimating the relative weighting of reward and likelihood on observed reach biases, we also examined the relationship between the observed reach biases and the relative subjective values of the targets in conditions when the rewards and likelihoods of the two targets were unequal. This relationship was examined both for direct reaches (A) and intermediate reaches (B). In both cases, the left panels show the distribution of power law fits to the relationship between reward ratio and the measured bias divided by the frequency ratio of the two targets (See Supplemental Methods). The right panel shows a histogram of the estimated value of the exponent (alpha) applied to the reward ratio; larger alpha values reflect less discounted reward relative to likelihood in determining the relative subjective values of the two targets.

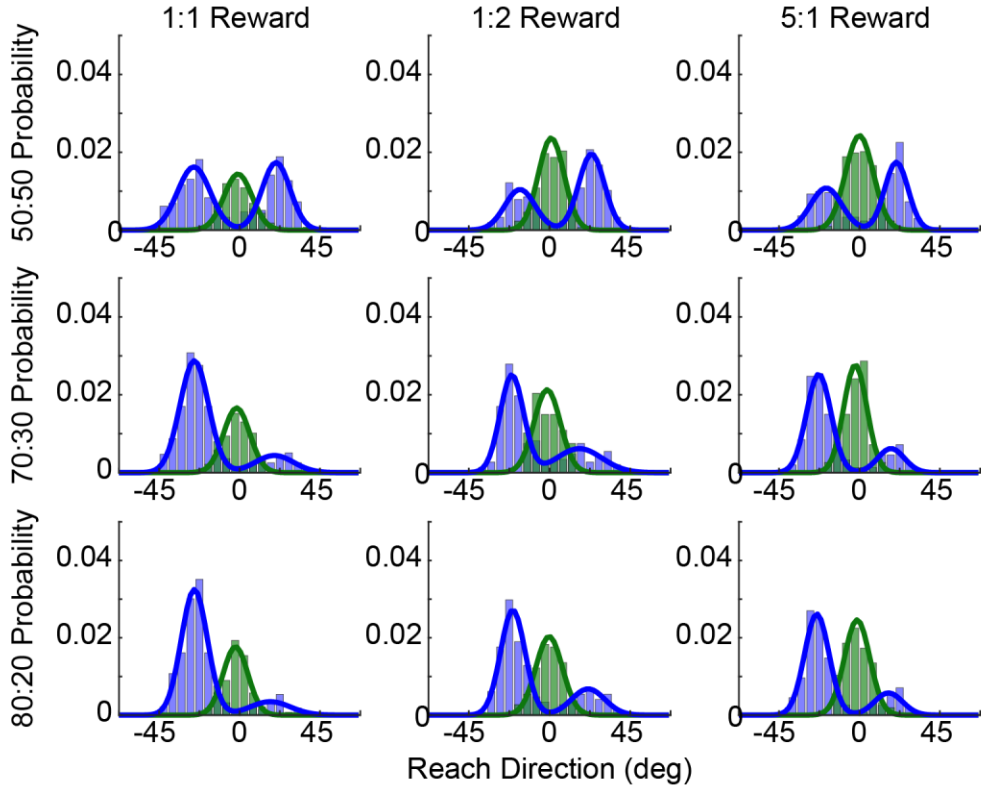


**Supplemental Figure 2.** Reach-direction histograms for Experiment 2. Initial reach direction for direct and intermediate reaches (data pooled across all participants), with varying reward and prospective-probability ratios. In all cases, the data have been aligned such that reaches to the more rewarded target are in the positive direction (rightward), and reaches to the more probable target are in the negative direction (leftward). As the relative reward ratio increases (left to right), reaches are biased toward the more rewarded target. Likewise as the relative prospective-probability ratio increases (top to bottom), reaches are biased toward the more likely target. Fits reflect the average distribution parameters from bootstrapped fits to subsets of the pooled group distributions.

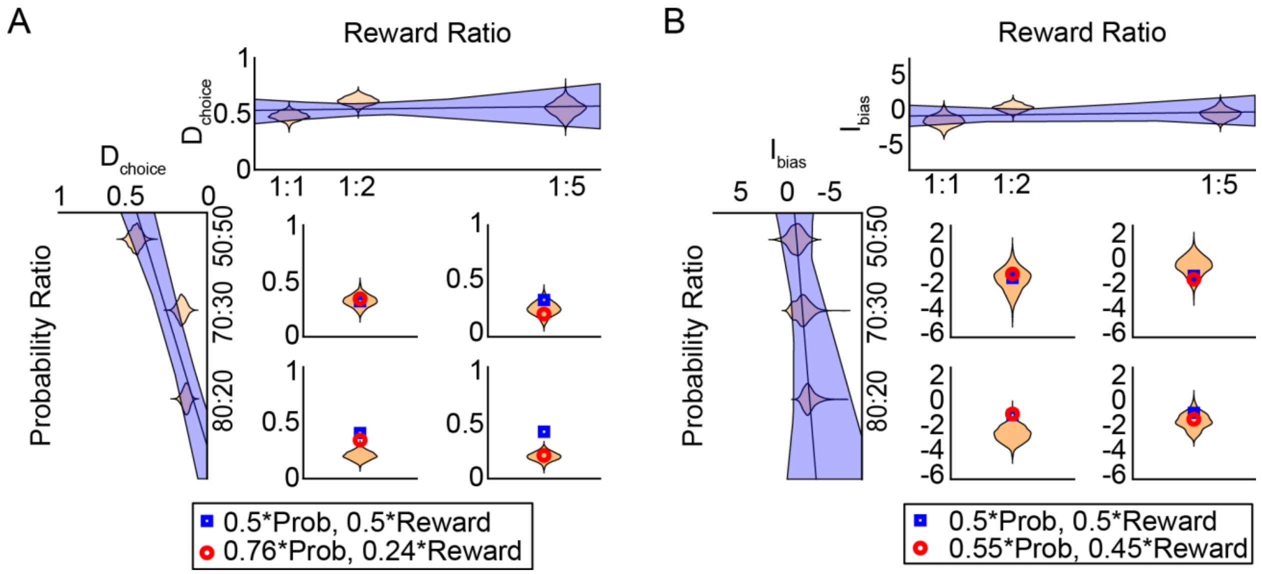


**Supplemental Figure 3.** Estimation of biases in Experiment 2. For direct reaches (A) and intermediate reaches (B), the effect of changing reward (top) or prospective-probability(left side) alone was fit by a linear regression. These regressions were then use to predict the reach direction bias observed when both reward and probability were changing (four small panels). Weighting the influence of probability and reward equally fit the data moderately well (blue squares); however, when the weights were allowed to vary we obtained much better estimates (red circles). These weights indicated a much stronger influence of probability compared to reward for direct reaches, but the influence of probability and reward were weighted more similarly for intermediate reaches.

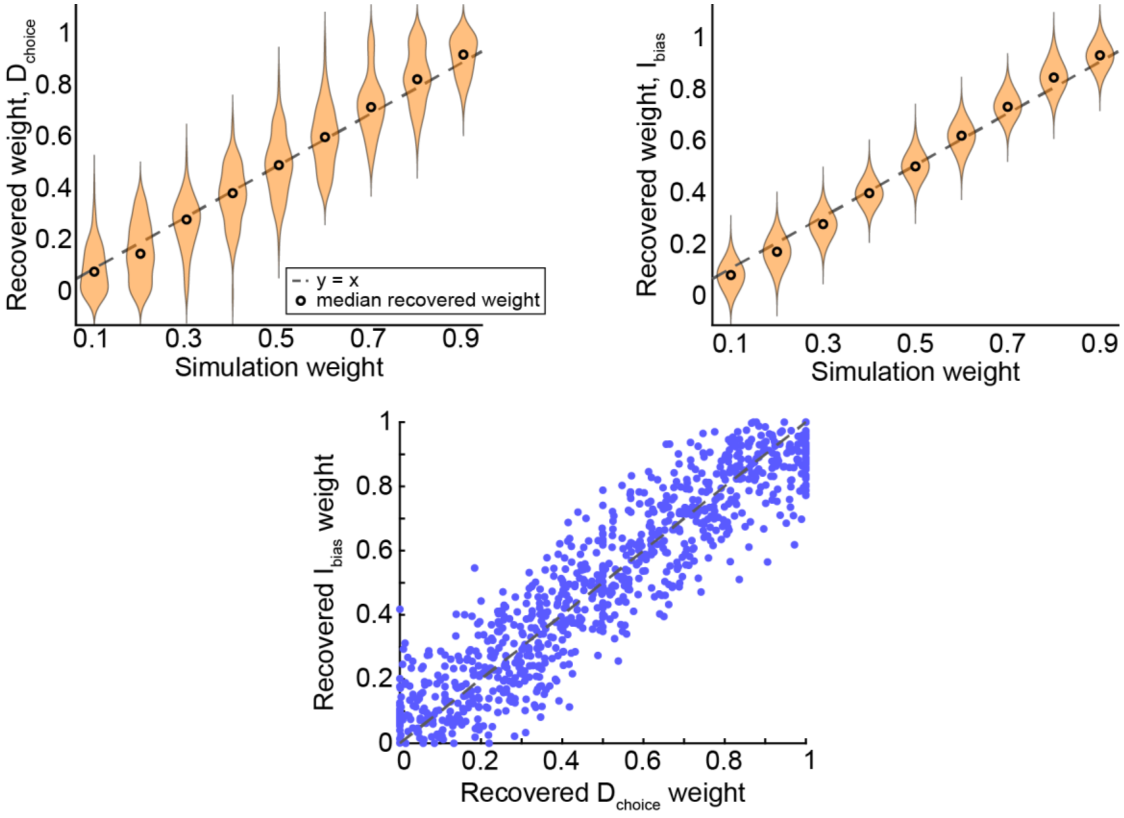


**Supplemental Figure 4.** Parameter recovery analysis to evaluate the weight-estimation method. In Experiment 3, weights for direct and intermediate reaches were highly uncorrelated. To examine whether this finding arose because the weights were actually uncorrelated or because the method used to estimate the weights was unreliable, we performed a parameter recovery analysis (see Supplemental Methods). We observed good recovery of our simulation weights (compare the median recovered weight to the simulated weight, black open circles), for both (A) direct reaches, and (B) intermediate reaches. The dashed gray line reflects the line y = x; all median recovered weight values fall close to this line. (C) We also compared the recovered weights from direct and intermediate reaches to each other on each simulation repetition, where in each simulation the same weight value was used to generate both direct and intermediate reaches. Note that these points cluster also along the line y = x, suggesting that if we see a lack of relationship between the weights estimated on direct and intermediate reaches across participants, it is likely because these weights were indeed uncorrelated rather than because of an artifact introduced by the weight-estimation procedure.

### SUPPLEMENTAL METHODS

#### Subjective Value Analysis

Typical neuroeconomic analyses model the subjective value ($SV$) of option $i$ as a power law relationship, typically assuming that the target reward ($R$) is discounted relative to the target likelihood ($P$) by some power ($\alpha$):

$$(3) \qquad SV_i = P_i {R_i}^{\alpha}$$

Here, we assume that the reach bias observed is proportional to the ratio of the subjective values of the two targets:

$$(4) \qquad \begin{aligned} Bias \propto \frac{SV_1}{SV_2} &= \frac{P_1 {R_1}^{\alpha}}{P_2 {R_2}^{\alpha}} \\ &= \left(\frac{P_1}{P_2}\right)\left(\frac{R_1}{R_2}\right)^{\alpha} \end{aligned}$$

To be able to directly compare the biases for direct and intermediate reaches in this analysis, we scaled the bias on intermediate reaches (i.e., the initial reach direction) to be on the same range as direct reaches (between 0 and 1, since bias is measured as the proportion of reaches to the more rewarding target). Thus initial reach direction on intermediate reaches was divided by the angular distance between the two targets (45°) and recentered about 0.5.

Once the biases were rescaled, we then estimated $\alpha$ for both direct and intermediate reaches, for each iteration of our bootstrapped data. This was done by fitting a power law to the relationship between the reward ratio of the two targets and the bias divided by the likelihood ratio of the two targets as in Equation 4 (see Supplemental Figure 1). Larger $\alpha$ values reflect less devaluation of target reward information relative to target likelihood information; or in other words, larger $\alpha$ values suggest that reward information exerts a stronger influence in determining the subjective value of the options, and hence more strongly influences the observed behavioral bias.

#### Parameter Recovery Analysis

To examine how realistically our weight-estimation method could accurately recover the underlying weights placed on frequency and reward biases, we performed a parameter recovery analysis. Biases were simulated in reach preference (direct reaches, $D_{choice}$) and reach direction (intermediate reaches, $I_{bias}$). These biases were determined by choosing a relative weighting of likelihood and reward. On each simulated trial, we calculated the preference for choosing the more frequent target based on the current trial's frequency ratio and the reward ratio separately (e.g., if the frequency ratio is 2:1, the preference for choosing the more frequent target is 0.67). We calculated the distance this preference was from equal preference (i.e., 0.5), and scaled that distance by the predetermined frequency or reward weighting; this yielded a frequency-biased and reward-biased preference for each target:

$$Target_1\_preference_{frequency} = \left(\frac{frequency_{Target1}}{frequency_{Target1}+frequency_{Target2}} - 0.5\right) * frequency\_weight + 0.5$$

$$Target_1\_preference_{reward} = \left(\frac{reward_{Target1}}{reward_{Target1}+reward_{Target2}} - 0.5\right) * reward\_weight + 0.5$$

Then, the objective utility (probability times reward) was computed for each of the two targets based on the scaled preference values:

$$EV[Target_1] = Target_1\_preference_{frequency} * Target_1\_preference_{reward}$$

$$EV[Target_2] = Target_2\_preference_{frequency} * Target_2\_preference_{reward}$$

These expected values were normalized by the sum of the expected values for the two targets to yield the probability of choosing each target. On direct reaches, a random number was drawn; if the random number was lower than the probability of choosing Target 1, we recorded a direct-reach choice of Target 1; otherwise we recorded a  choice of Target 2. On intermediate reaches, we converted the probability of choosing Target 1 into a reach direction by multiplying by 30°, then subtracting 15° to center reaches about the midline. In this way, we could build a binary distribution of direct reaches and a continuous distribution of intermediate reaches, for every possible combination of frequency and reward ratios presented during Experiment 3 (200 trials were simulated for each condition). Finally, we applied our weight estimation approach as described in our Methods (computing regressions to estimate the relationship between frequency ratio and bias and reward ratio and bias separately, then finding the best weighting of these regressions to explain the bias observed when both frequency and reward ratios were not equal to 1) to recover the degree to which frequency was weighted across simulated trials ($D_{choice}$ and $I_{bias}$). We repeated this simulation for 100 iterations at each of 9 frequency-reward weightings (0.1 to 0.9). We compared the recovered weighting to the actual weighting for both direct and intermediate reaches separately (Supplemental Figure 4), as well as comparing the recovered weightings for direct reaches versus intermediate reaches (since on each repetition both direct and intermediate reaches were simulated from the same underlying weights).
